# Stimulated emission depletion microscopy with a single depletion laser using five fluorochromes and fluorescence lifetime phasor separation

**DOI:** 10.1101/2022.04.13.487856

**Authors:** Mariano Gonzalez Pisfil, Iliya Nadelson, Brigitte Bergner, Sonja Rottmeier, Andreas W. Thomae, Steffen Dietzel

## Abstract

Stimulated emission depletion (STED) microscopy achieves super-resolution by exciting a diffraction-limited volume and then suppressing fluorescence in its outer parts by depletion. Multiple depletion lasers may introduce misalignment and bleaching. Hence, a single depletion wavelength is preferable for multi-color analyses. However, this limits the number of usable spectral channels. Using cultured cells, common staining protocols, and commercially available fluorochromes and microscopes we exploit that the number of fluorochromes in STED or confocal microscopy can be increased by phasor based fluorescence lifetime separation of two dyes with similar emission spectra but different fluorescent lifetimes. In our multi-color FLIM-STED approach two fluorochromes in the near red (exc. 594 nm, em. 600-630) and two in the far red channel (633/641-680), supplemented by a single further redshifted fluorochrome (670/701-750) were depleted with 775 nm. To the best of our knowledge, these are the first published five color STED images. Generally, this approach doubles the number of fully distinguishable colors in laser scanning microscopy. We provide evidence that eight color FLIM-STED with a single depletion laser would be possible if suitable fluorochromes were identified and we confirm that a fluorochrome may have different lifetimes depending on the molecules to which it is coupled.

## Introduction

In the life sciences, highly resolved images of a single structure can be helpful in some cases, but often it is necessary to know where one structure is in relation to another or several others. This generates a demand for multi-color super-resolution microscopy. Commercially available stimulated emission depletion (STED) microscopes have up to three depletion laser wavelengths, 592 or 595, 660 and 775 nm. This theoretically allows usage of fluorochromes over the full spectral range of a confocal microscope. However, in the experience of others^1^ and ourselves, dyes depleted with a 592 or 660 nm continuous wave laser tend to bleach fast, compared to some that can be depleted with the pulsed 775 nm laser. This is also reflected by the relatively small number of fluorochromes typically used in STED microscopy (see following examples). Further technical obstacles for usage of several depletion lasers are the need to align those lasers very precisely to allow meaningful super-resolution multi-color colocalization studies and total photobleaching of long wavelength dyes by the lower wavelength depletion laser. Therefore, ideally only one depletion laser is used for all fluorochromes^2^. And indeed, multi-color STED studies tend to apply only 775 nm depletion using near or far red dyes^1-8^.

To our knowledge, the highest number of distinguishable fluorochromes used in a STED publication so far was four^5,9^. Rönnlund et al. applied a scheme with two color channels where two dyes or distinguishable dye conjugates were in each color channel (ATTO 647N & Dylight650; Alexa Fluor 594 coupled to antibody and phalloidin). After an initial scan, one in each channel was specifically photobleached. While this was feasible in small and flat platelets (thrombocytes), it may be difficult in larger cells, e.g. due to sample drift. The 2017 study from the group of Stefan Hell, who invented STED microscopy, excited four dyes with 612 nm and depleted with 775 nm in fixed cells. They used the fluorochromes ATTO 594 (max exc./em 601/627), Abberior Star 635P (633/654), KK1441 (661/679), and CF680R (680/701)^5^, dyes with heavily overlapping emission spectra which they separated computationally by mathematical spectral unmixing. The anti-stokes excitation^10,11^ of CF680R by the STED beam was subtracted prior to the unmixing. A ‘residual crosstalk’ between channels of up to 20% remained in a selected image. Unfortunately, one of these dyes, KK1441, does not seem to be commercially available, neither easily replaceable, thus limiting the number of usable dyes from this scheme for most researchers to three. Both groups performed their experiments on custom-built microscope systems, indicating a talent and preference for microscope building, which in the case of the average life scientist who wants to examine biological questions is usually not given.

A further approach to multi-color STED used up to four primary antibodies with single strand DNA labels (∼ 10 nucleotides) to which complementary DNA oligonucelotides bound transiently^12,13^. One of the four complementary DNA strands was perfused over the sample and washed out before the next strand was applied. This approach theoretically allows unlimited numbers of labels. Potential problems are z-drift or shift between recordings (in particular if several fields of view were to be recorded), the long time periods needed for each image (about 20 min per color in fast variants^14^), and the inability to make permanent samples. The need to conjugate the DNA oligonucelotides to the primary antibodies and the impossibility to preselect multiple labeled areas of interest through the eyepiece or with an overview scan may further deter life scientists from this approach.

A combination of STED with fluorescence lifetime imaging (FLIM) to separate fluorochromes was first published in 2011, also by the group of Stefan Hell^2^. They separated ATTO 647N and KK 114^6^ (later commercialized as Abberior Star Red, according to the company’s web site) by lifetime and used ATTO 590 as a third color with a second STED beam. Since with FLIM separated fluorochromes the photons of two dyes are recorded simultaneously, any drift or misalignment issues are excluded by design. However, a decade ago FLIM equipment was slow and difficult to operate compared to a confocal microscope. In addition, classical curve fitting analysis to separate dyes by lifetime requires high numbers of photons^15^, which are typically difficult to achieve in STED microscopy of biological samples. Curve fitting for dye separation faces another problem: While many dyes may exhibit a single exponential lifetime free in solution, when coupled to antibodies or other targeting molecules they can obtain additional lifetime components. Two fluorochromes with bi-exponential behavior would thus present four lifetime components, rendering classic exponential decay fitting more complex^15^. With increased number of lifetime components the number of required photons and thus imaging time increases, making this approach difficult to combine with STED.

Accordingly, we found only three other papers, one of them also from the Hell group^16^, describing the use of FLIM to increase the number of fluorochromes in multi-color STED. In confocal microscopy, separation of five fluorochromes excited by the same laser line was demonstrated by a mixture of spectral and FLIM separation^15^. Another study made use of cross labelling antibodies to create new lifetime species via Förster resonance energy transfer (FRET), later unmixed by the same spectral FLIM separation approach^17^. However, for high precision color separation a commercially not available spectral detector was used in both studies and the applied pattern matching for color separation requires relatively high photon numbers, at least if structures are overlapping^15,18^. The method was also applied to STED, but here the separation of only two dyes in a single color channel was demonstrated on images with 5,000 - 10,000 photons in the brightest pixels^15^.

Complementing fluorescence lifetime and emission spectra, utilization of the absorption spectra was suggested for multi-color STED^18^, but so far the number of fluorochromes was limited to three.

Recent advances in FLIM technology brought faster systems which are fully integrated with commercial confocal and STED software and thus much more user friendly and easier to operate by life scientists. In addition, the development of phasor based analysis of FLIM data^19-21^ reached a point where it allows easy and fast fluorochrome separation with substantially lower photon numbers when compared to curve fitting if background is present^22^. Applicability with low photon numbers is important since in STED, due to the depletion of fluorochromes and much smaller pixels compared to typical confocal microscopy, in a given time span the number of detected photons per pixel is low. Phasor analysis with low photon numbers will not determine absolute lifetimes with the same accuracy as curve fitting with high photon numbers. However, to separate fluorochromes only relative lifetime differences between two dyes are needed. We therefore revisited the multi-color FLIM-STED approach and combined it with phasor analysis.

We used a commercial confocal system with STED and FLIM capabilities to develop a multi-color FLIM-STED scheme with one depletion laser (pulsed 775 nm) and one multi-wavelength excitation laser, thus minimizing alignment issues. Only commercially available fluorochrome-labeled antibodies, phalloidin and wheat germ agglutinin (WGA) were used. Fixation and staining were performed by typical immunofluorescence protocols and are thus easily reproducible by other life scientists. We show for biological samples that phasor based image analysis can very well separate two fluorochromes by lifetime in a spectral channel. We demonstrate five color STED and five color confocal images in only three spectral channels. We also suggest avenues to further increase the number of colors for STED with a single depletion laser.

## Results

### Five color STED microscopy

We performed FLIM-STED with five fluorochromes using one depletion laser and three spectrally separated channels, each with a fitting excitation wavelength. Two of the channels carried two spectrally similar fluorochromes that were separated by phasor lifetime analysis (Figure 1). Fixed HeLa cells were stained in the far red channel with Abberior Star 635P coupled to a secondary anti-rabbit antibody (Tau 2.0 ns) to detect mitochondrial outer membrane protein TOMM20 and with ATTO 647N coupled to phalloidin (Tau 3.5 ns) which delineates the actin cytoskeleton. Excitation was at 633 nm, detection at 641-680 nm. All lifetimes are given for confocal images.

**Figure 1:**
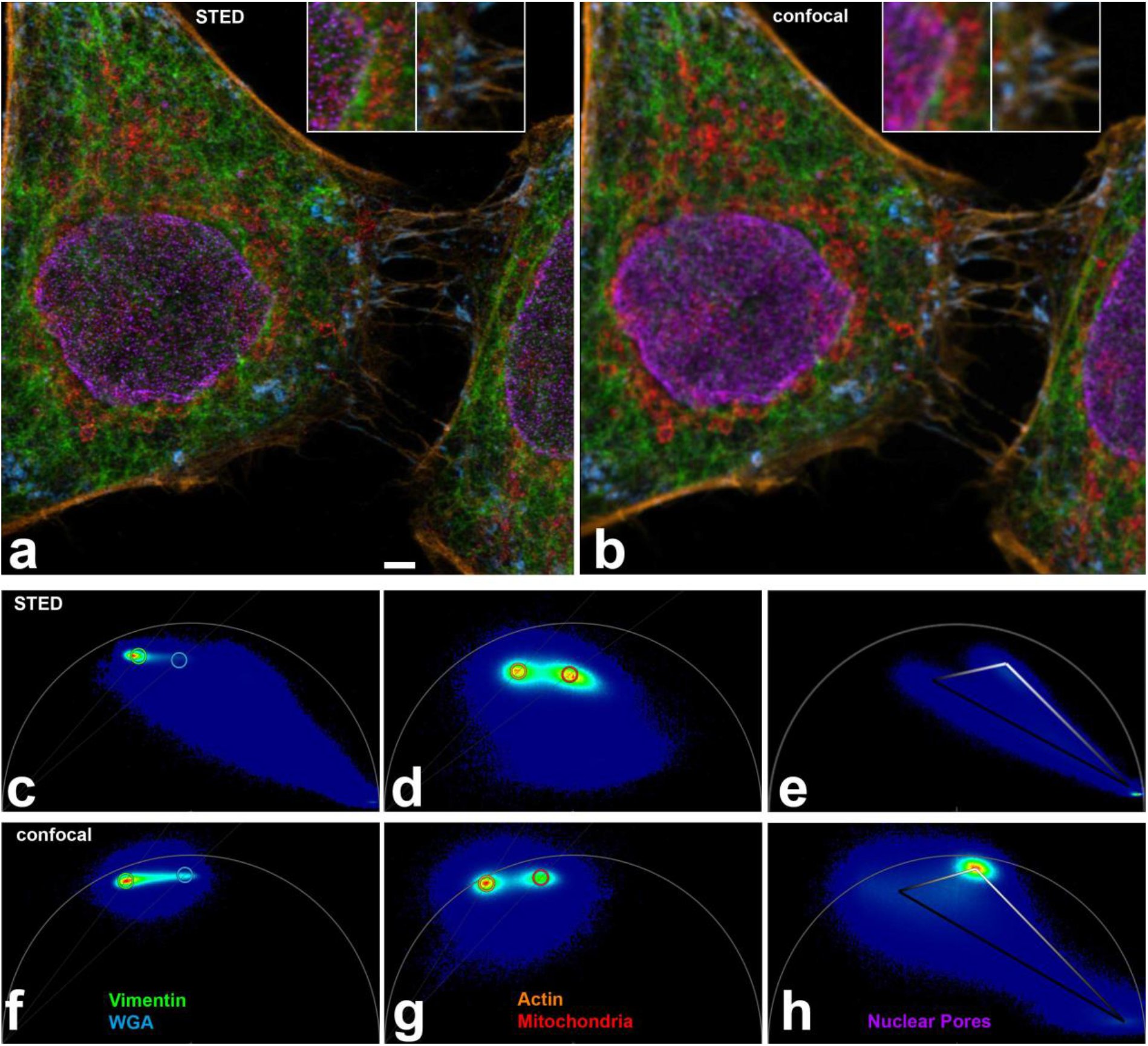
(a) Five color FLIM STED image. Insets show two areas doubled in size. Raw images had 1516, 1213, and 542 photons in the brightest pixels in the near red, far red and CF680R channels. Scale bar 2 µm. (b) Corresponding FLIM confocal image with the same pixel size (25 nm). (c-h) Phasor plots from STED (c-e) and confocal (f-h) raw images corresponding to (a,b). Near red channel (c,f, 600-630 nm), far red channel (d, g, 641-680nm) and 701-750 nm channel (e,h) are shown. In phasor plots, each image pixel is positioned according to its lifetime behavior. Pixels with monoexponential decay will be on the semi-circle (called ‘universal circle’), those with multiexponential decay towards the inside. Short lifetimes are to the right, long lifetimes to the left, the scale is non-linear. Note that the lifetime distribution in phasor plots in STED is shifted to the right, reflecting shorter lifetimes. For nuclear pores, the confocal phasor plot (h) shows pixels with crosstalk from the far red channel (compare g) while the STED phasor plot (e) shows in addition strong reflection of the 670 nm excitation in the lower right corner (zero lifetime), since no notch filter was available to suppress it.

The near red color channel (594 nm, 600-630 nm) was populated with Alexa Fluor 594 coupled to a secondary anti-chicken antibody (Tau 3.5 ns) to visualize the cytoskeletal filament vimentin and WGA-CF594 (Tau 1.9 ns). WGA is a lectin that has a high affinity for sialic acid and N-acetylglucosamine moieties of glycoproteins and glycolipids; thus it stains the plasma membrane^23-25^.

In the third spectral channel, CF680R labeled a secondary anti-mouse antibody (Tau 1.2 ns) to detect nuclear pores and was excited with the longest available wavelength, 670 nm (detection 701-750 nm). Sequential scanning of the three channels allowed to minimize bleed-through of fluorochromes to neighboring spectral channels and to adapt the STED laser power. By the latter, we could avoid the strong anti-stokes excitation of CF680R by the 775 nm STED beam that was previously described^5^ (Figure 1a,b).

Phasor based lifetime separation of two fluorochromes in the same color channel in STED images (Figure 1c-e) revealed excellent partitioning (Figure 1a). Visual comparison with confocal images of likewise acquisition (Figure 1b, f-h) revealed significant resolution improvement.

In raw images of the CF680R channel, it was obvious that a substantial cross talk from ATTO 647N and to a much lesser extent from Abberior Star 635P was present, when excitation was performed with 670 nm (Figure 2a). In our routine procedure, this unwanted signal was efficiently eliminated by a Tau-STED approach implemented in the software, making use of the large differences in fluorescence lifetimes (1.2 vs 2.0 and 3.5 ns). For testing purposes, we could generate a few images on a microscope that allowed longer excitation wavelengths. With 685 nm excitation, clear raw images of nuclear pores were obtained without noticeable ATTO647N or Abberior Star 635P contribution (Figure 2b-d).

**Figure 2:**
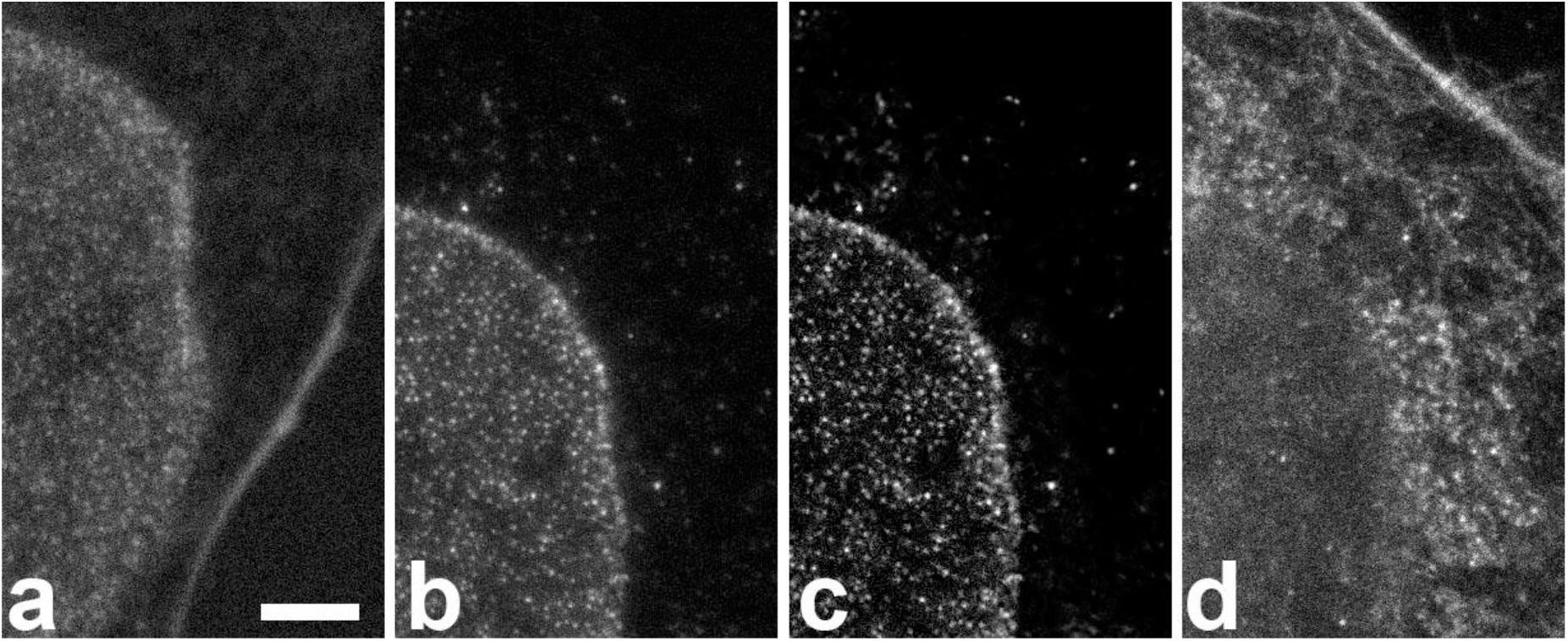
Crosstalk of far red dyes into the CF680R nuclear pore channel is observed with 670 nm excitation but not with 685 nm. Scale bar 2 µm for all images. (a) Excitation of a five color sample with 670 nm. The detection window was 701-750 nm. This raw image prior to Tau-STED processing shows substantial crosstalk from ATTO 647N (actin staining, strong fiber in the image from bottom to right) while Abberior 635P (mitochondria) crosstalk is hardly above background. (b) Excitation of a different cell with 685 nm on another microscope shows no sign of crosstalk from other dyes. A haze around the nuclear pores is due to anti-stokes excitation of CF680R by the depletion beam. (c) The same image after Tau-STED processing which removed the anti-Stokes contribution. (d) Raw image of the area shown in b, c in the far red channel with actin and mitochondrial staining, demonstrating the presence of both structures and their fluorochromes.

### Orange as an additional spectral channel

A new probe for the orange channel (exc. ∼550, em. 560-600), SPY555-actin, for which the manufacturer’s material suggested that it could be depletable with 775 nm, became available during the course of this study. This was indeed the case in our experiments (Figure 3). The super-resolution effect was clearly detectable, suggesting that this spectral range could be used as a fourth spectral channel in STED with 775 nm depletion, with two additional dyes, separable from each other by lifetime, if suitable fluorochromes were available.

**Figure 3:**
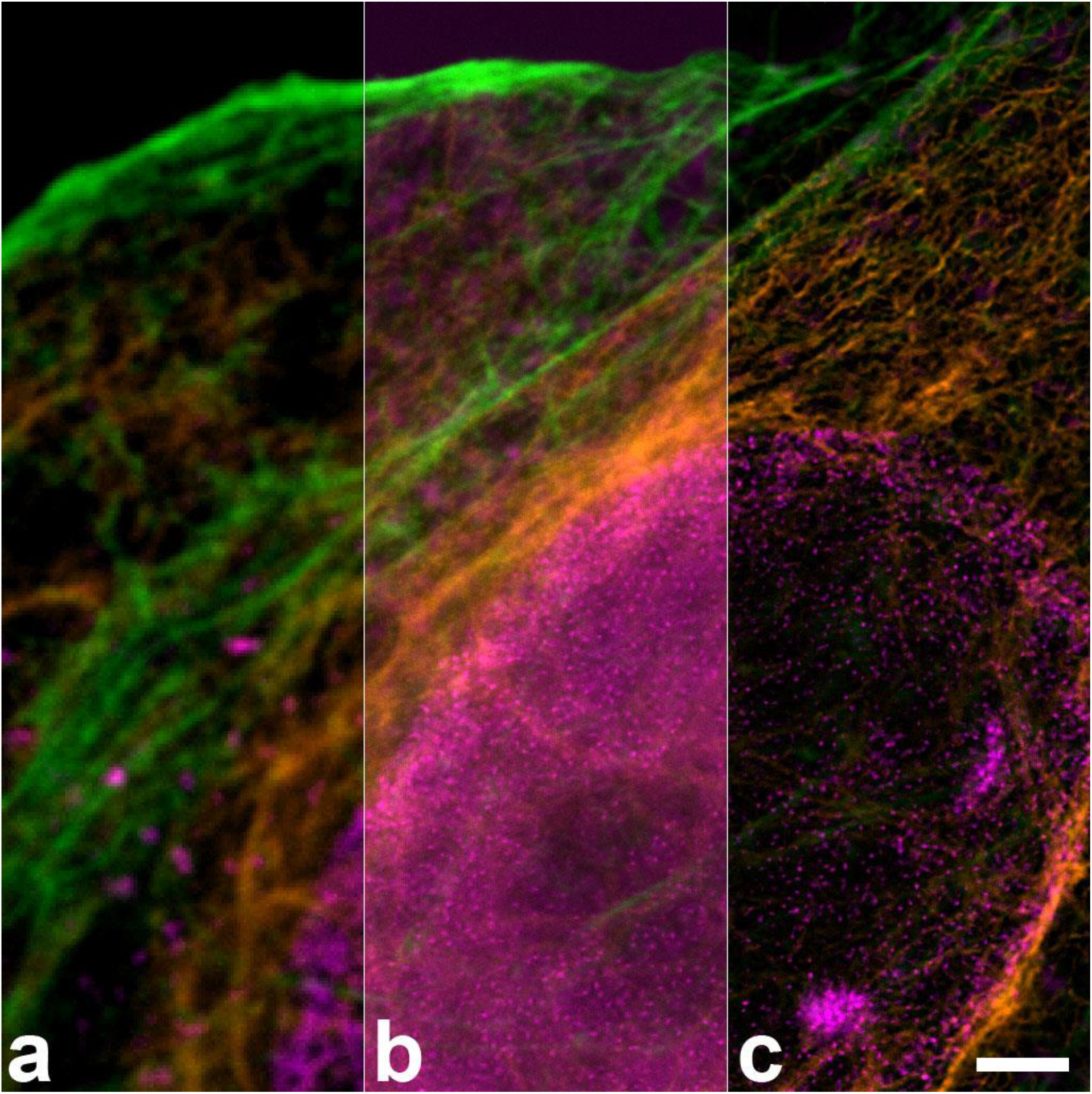
Three color image with spectrally separated SPY555-actin (green) and antibody stainings against vimentin (Alexa Fluor 594, orange) and nuclear pores (CF680R, magenta), all depleted with 775 nm. (a) confocal, (b) STED (c) Tau-STED. This dye combination could be supplemented with e.g. Abberior Star 635P to obtain four spectrally separable fluorochromes. Scale bar: 2 µm.

### STED causes a shortening of fluorescence lifetimes compared to confocal images

When comparing phasor plots of STED and confocal images, it was obvious that the average lifetimes of all dyes were substantially shortened in STED, compared to confocal images (Figure 1c-h). Fluorochrome molecules that stay longer in the excited state are more likely to undergo stimulated depletion while those with a very short residence time in the excited state may have already emitted a short-lifetime fluorescence photon before stimulated depletion occurred^26,27^. Accordingly, we too observed that lifetime was inversely related to STED laser power.

The shortening of the lifetimes makes phasor separation for STED images (Figure 1c-e) more demanding than for confocal images (Figure 1f-h), since the absolute difference between lifetimes of two fluorochromes also becomes smaller. The lifetime of a fluorochrome in STED depends on its depletability by the wavelength of the STED beam and the beam’s intensity and shape. STED lifetime in any specific experiment is thus difficult to predict. This suggests a trial-and-error approach to identify new dye pairs in the same color channel suitable for FLIM-STED, starting with fluorochromes that are reported to have big lifetime differences and that show low photobleaching in STED conditions.

### Two colors with a single fluorochrome: lifetime depends on the binding partner

When testing various combinations of stainings for this study, a combination of two antibodies labeled with either Abberior Star 635P or ATTO 647N showed more similar lifetimes for both dyes compared to the experiment presented above where ATTO 647N was coupled to phalloidin. In STED images, the two antibody conjugated dyes could not be separated sufficiently by the phasor approach.

Different lifetimes of these two dyes and others, depending on their conjugation partner, were described previously^2,15^. In our hands, the lifetime differences induced by different conjugation partners (antibody or phalloidin) of Abberior Star 635P or ATTO 647N were sufficient for discrimination of the two conjugates by phasor based separation in confocal images but did not lead to optimal results in STED, due to the generally shortened STED lifetimes. We performed double stainings with the same fluorochrome (either Abberior Star 635P or ATTO 647N) coupled to both, an antibody and to phalloidin (Figure 4). For both dyes, phasor analysis of confocal images allowed separation of phalloidin and antibody conjugates. Abberior Star 635P coupled to phalloidin had a lifetime of 3.2 ns, but only 2.3 ns when coupled to an anti-chicken antibody to detect vimentin (Figure 4a). Lifetime for ATTO 647N was measured as 3.6 ns when coupled to phalloidin but as 2.8 ns when coupled to an anti-mouse antibody to detect nuclear pores (Figure 4b). Thus in both cases, the phalloidin coupled dye had a 0.7 – 0.9 ns longer lifetime than the antibody coupled one, a sufficient difference for phasor based separation in confocal images (Figure 4c,d). In STED images, however, the lifetimes and the difference between them decreased so that a phasor based separation did not achieve a satisfying quality.

**Figure 4:**
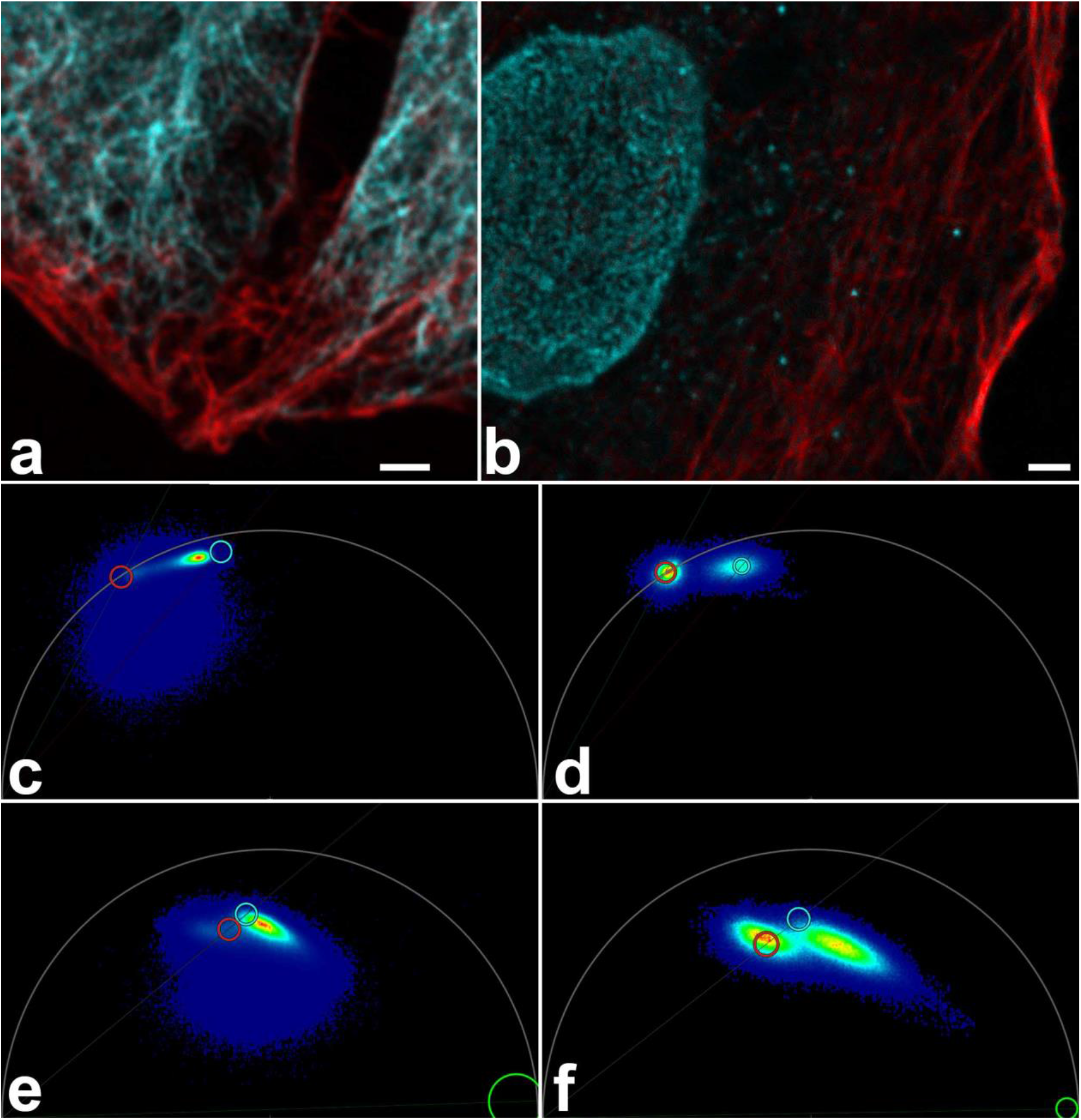
Fluorochromes have different lifetimes when coupled to antibodies or to phalloidin. (a) Confocal image of Abberior Star 635 P coupled to an antibody delineating vimentin (cyan) and to phalloidin (actin staining; red) and subjected to phasor based lifetime separation. (b) Confocal image of ATTO 647N coupled to phalloidin (actin staining, red) and an antibody (nuclear pores, cyan. Both confocal images show good separation of the two stained structures. Scale bars 2 µm. (c,d) Phasor plots for the complete raw images of (a,b). The actin coupled dyes have longer lifetimes (left cloud, red circle) compared to their antibody-conjugated versions (right cloud, cyan circle). (e,f) Phasor plots of respective STED images show that the lifetime differences are much reduced. Therefore, separation of STED images was not satisfying and resulting images are not shown. Circles indicate the best possible positions for separation of the two species. In e and f the right-hand clouds contain the pixels with antibody label derived photons, but they also contain substantial amounts of actin label derived photons. To compensate for this the circles were positioned next to the clouds.

### Three fluorochromes were not satisfactory discriminated by phasor plot separation

In an attempt to further increase the number of fluorochromes per color channel, we performed a triple staining with two secondary antibodies labeled with either Abberior Star 635 visualizing vimentin or with Alexa Fluor Plus 647 visualizing mitochondrial outer membrane, while the actin cytoskeleton was delineated by phalloidin-ATTO 647N (Figure 5a). Vimentin, mitochondria and actin filaments form longitudinal structures that cross each other within a cell. Although the lifetimes of these dyes were quite different (Tau = 2.0, 1.2, and 3.5 ns, respectively), already in confocal images it turned out that three dyes could not be separated satisfactory by our current phasor approach (Figure 5b,c). While some components were split up correctly into the respective channels, finer details got lost and possibly misallocated. For many image pixels it was impossible to distinguish whether their signals and average lifetime were made up by the dye with the intermediate lifetime, or by a mixture of the long and short lifetime dyes or both (Figure 5b).

**Figure 5:**
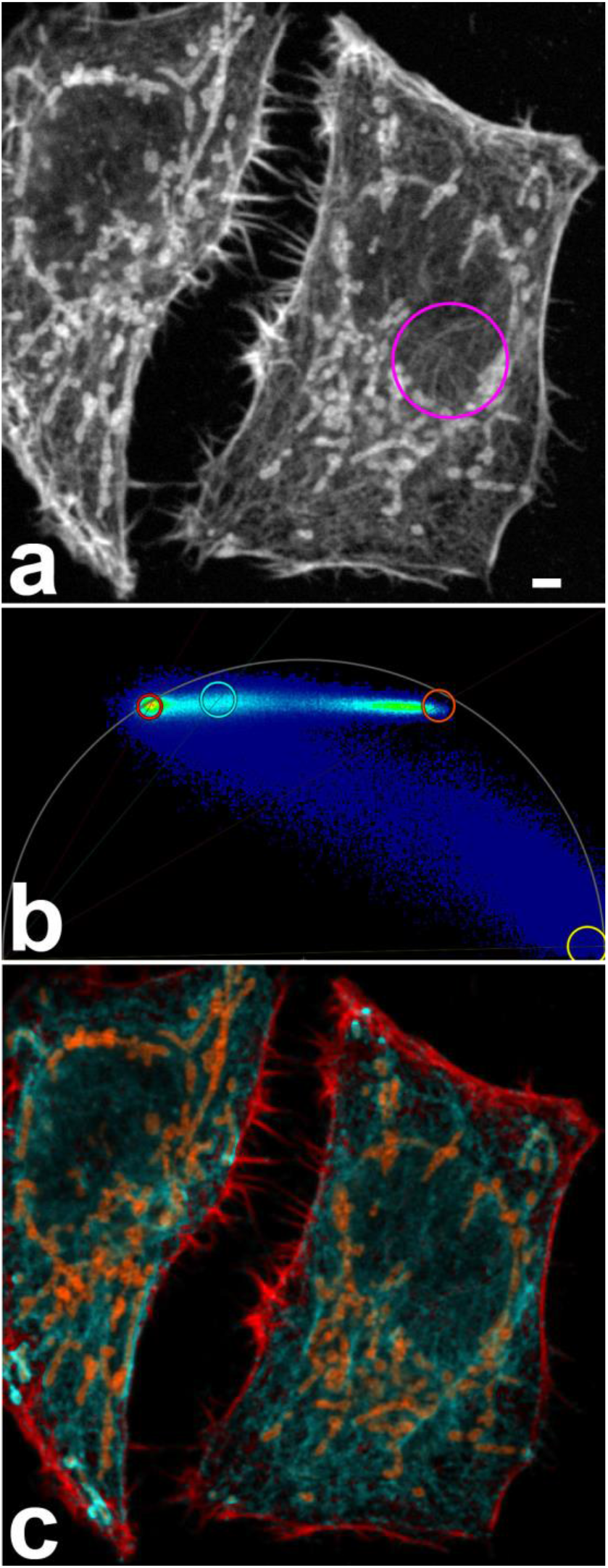
Three fluorochromes in the same spectral range with different lifetimes cannot be separated with high quality by the phasor approach. (a) Confocal raw image with Abberior Star 635P labeled vimentin, Alexa Fluor Plus 647 labeled mitochondria and ATTO 647N labeled actin. Scale bar 2 µm. Magenta circle highlights some delicate vimentin fibers. (b) Phasor plot of this image. The lifetime components are indicated by circles: ATTO 647N-actin red, AS635P-vimentin cyan, AF647-mitochondria orange. A background component (yellow circle) was also included for phasor separation to suppress noise. In pixels with a low photon count, noise from reflection and other sources will determine the average lifetime of the pixel, making it likely to be sorted with the background component. (c) Phasor separated image, same color code. Note the poor preservation of delicate vimentin fibers in the circle in (a). The respective pixels were not correctly assigned to the vimentin channel.

A separation of three (Figure 5c) or even more currently available dyes might be successfully performed when a certain amount of crosstalk can be tolerated, or if differently labeled structures do not overlap in space. However, it does not appear to be an advisable strategy for subcellular high resolution colocalization studies. In such studies, a successful separation of a third dye could potentially be possible if it had a very different pattern in the phasor plot, e.g. with a lifetime of over 10 ns. Then three fluorochromes might be arranged in a triangle in the phasor plot, not a line. However, typical commercial fluorochromes have a lifetime below 4 ns and are thus relatively close to each other in the phasor plot.

### 3D-stacks and fluorescence lifetime

We investigated whether our FLIM-STED approach can be applied to record 3D-stacks of images. We found that measured lifetimes change within the stack when moving from coverslip to deeper sample regions. A strong decrease of the average pixel lifetime towards the coverslip was found for CF680R-labeled nuclear pores, where we could not suppress reflection of the excitation beam with a suitable notch filter. The explanation that reflection was key in this case was supported by the observation that for CF680R-labeled nuclear pores the effect was also observed in confocal images. Since lifetimes were evaluated pixel-wise and reflected photons have a lifetime of zero, photons reflected at the coverslip surface shift the average lifetime of a pixel to smaller values.

No lifetime shift in confocal images towards the coverslip was observed for near red and far red dyes, where notch filters were available to suppress reflection. Still, in STED images lifetimes changed with distance to the coverslip (Figure 6). We assume that differences in the refractive index between coverslip and mounting medium and/or increasing refractive index mismatch with increasing depth induced shape changes of the depletion donut, thus causing the observed effect. The effect was large enough that phasor based lifetime separation would have to be performed for each z-position separately, rendering 3D recordings difficult.

**Figure 6:**
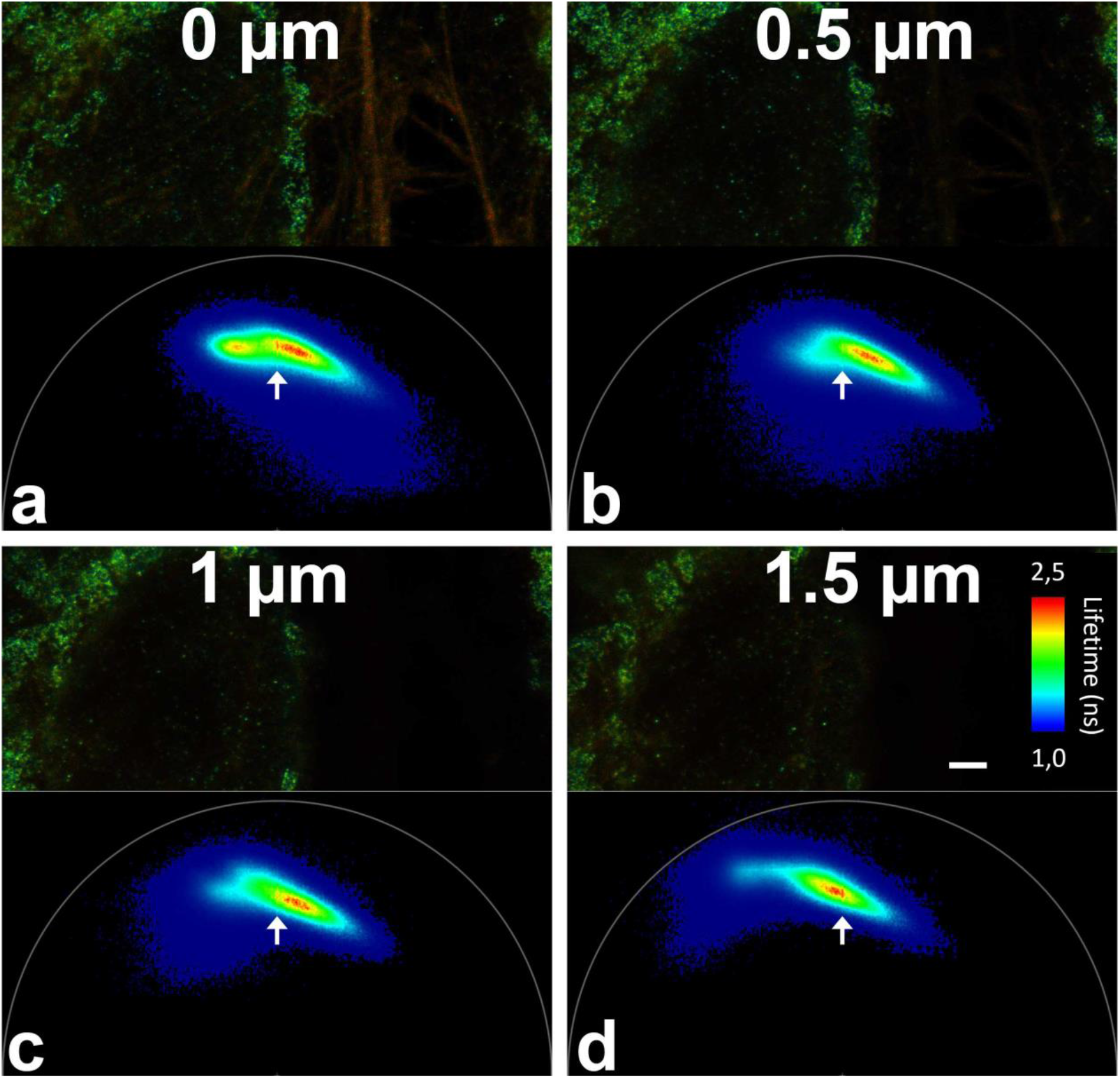
Fluorescence lifetime changes with distance to the coverslip. Parts of raw images of the far red channel from a five color sample at different sample depths (scale bar 2 µm) and their respective phasor plots. The color in the microscopic images codes the average lifetime of the pixels. (a) is near to the coverslip, (b-c) each 0.5 µm further into the sample. The arrows over the phasor plots always point to the same position. Near the coverslip, ATTO 647N labeled phalloidin produces a separate peak to the left in phasor plot that dwindles further into the sample. Note the movement of the Abberior Star 635P contribution in the phasor plot (major peak) relative to the arrow.

### Estimate of resolution improvement

In the five color STED images, all five labeled targets showed substantial resolution improvement when visually compared to confocal images (Figure 1). To obtain an estimate of resolution improvement, we applied Fourier ring correlation (FRC) to matching STED and confocal images. FRC works unsupervised and is thus unbiased by selection of individual structures for analysis. A disadvantage is that image areas with no or no small structures will generate a low resolution value and thus skew the average value of an image to higher, worse resolution values that do not represent the best resolution achieved in an image. FRC, however, does allow an unbiased comparison of two conditions such as confocal and STED. Five confocal and five STED five-color image sets were analyzed. The average FRC-calculated resolution values for confocal images in the five colors were between 276 and 305 nm. The corresponding average FRC values for STED were between 97 (WGA-CF594) and 136 nm (ATTO 647N, actin label), indicating a resolution improvement by a factor of ∼2.5.

### Crosstalk in 5 color FLIM-STED

For biological questions such as colocalization or non-colocalization of two proteins, it is important to assess how much of a given signal spills over to the neighboring channels and contaminates the obtained images, an effect known as crosstalk or bleed-through. In our study, ‘neighboring’ refers to spectral neighbors as well as to the two lifetime channels detected in one spectral channel. We compared two channels by evaluating photon numbers in a region of interest (ROI) that had strong signals in the ‘home’ channel but no structure of the neighboring channel was present, e.g. a strong vimentin signal in the vimentin channel (home) but no mitochondrial signal in the mitochondrial channel (neighbor). For the given example, the ROI was obtained by setting a high threshold to the vimentin signal but subtracting areas with high to low signals of mitochondria (low threshold; see Methods for details).

Sample preparations may have vastly different staining intensities in different channels. When comparing biological images, the spillover from neighbors should be set in relation to the home channel signal intensity to determine whether the spillover is problematic or not. To this regard, we calculated a “contribution” value in which bleed-through is normalized by the relative staining intensities of the signals in both channels (see Methods). Contribution may hence be reduced by changes to the staining procedure, such as different antibody concentrations or switching labels for individual cellular targets.

To determine which amount of crosstalk can be expected in purely spectrally separated color channels, we first analyzed a STED experiment with only three, spectrally separated fluorochromes: Alexa Fluor 594 (vimentin), Abberior Star 635P (mitochondria), and CF680R (nuclear pores) (Figure 7a). Eleven image sets with three images each were evaluated (Figure 7b). The average contribution was up to around 20%.

**Figure 7:**
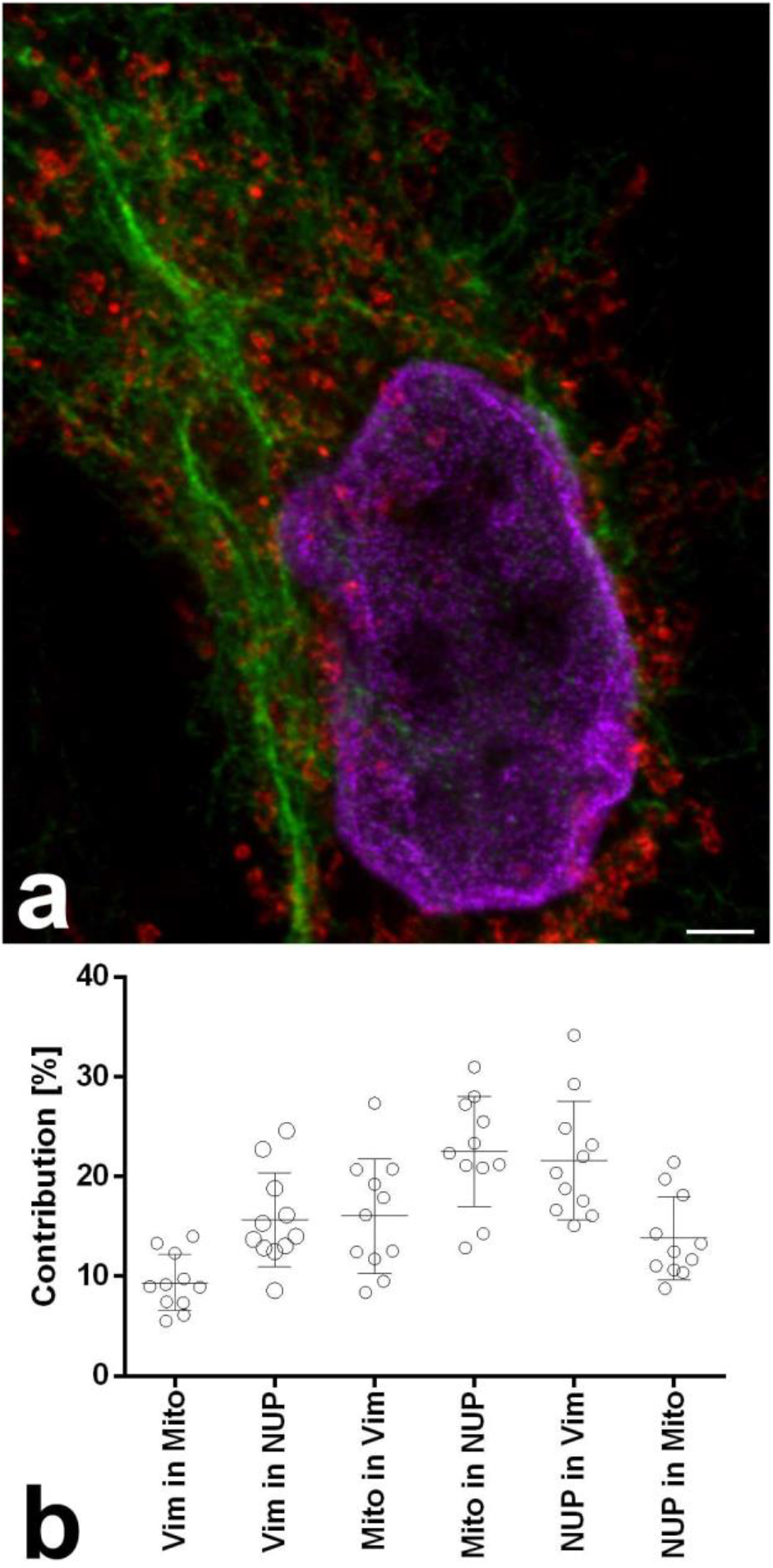
STED with three spectrally separated colors and corresponding crosstalk to other channels. (a) Example color overlay of the three channels. Vimentin Alexa Fluor 594 (green), mitochondria Abberior Star 635P (red), and nuclear pores CF680R (magenta). Scale bar 2 µm. (b) Contribution chart, n=11 image sets.

This evaluation also shows a clear limitation of the cross-talk determination in multi-color samples: The second highest value of 21.6% contribution was obtained for CF680R-labeled nuclear pore spillover into the Alexa Fluor 594-labeled vimentin channel. Clearly, strong spillover of CF680R into the near red channel is physically impossible and the observed phenomenon is due to biological overlap of the two structures. The brightest nuclear pore signal at the edge of the nuclei is in close proximity of a strong component of the vimentin around the nucleus. Despite the measures described above, such effects cannot be excluded in biological samples. All values are therefore an upper estimate. Actual crosstalk will be equal or less to the obtained values. Other average contribution values ranged from 9.4% to 22.5% (Figure 7b).

In five color FLIM-STED images described above (Figure 1a), average contribution values from 21 image sets were between 1.7 and 14.9% (Figure 8). Notably, the distributions of the four contributions between lifetime separated channels from the same spectral channel (Abberior Star 635P-labeled mitochondria versus ATTO647N-labeled phalloidin-actin and CF594-labeled WGA staining versus AlexaFluor594 labeled vimentin) were well within the range for contributions between spectrally separated channels (Figure 8). Again, the highest contribution mean value was obtained for CF680R-labeled nuclear pore contribution into AlexaFluor594-labeled vimentin. As above, we contribute this solely to biological overlap, not to physical spillover of photons to the near red channel. We conclude that crosstalk between lifetime separated images is well within the range of crosstalk between spectrally separated channels.

**Figure 8:**
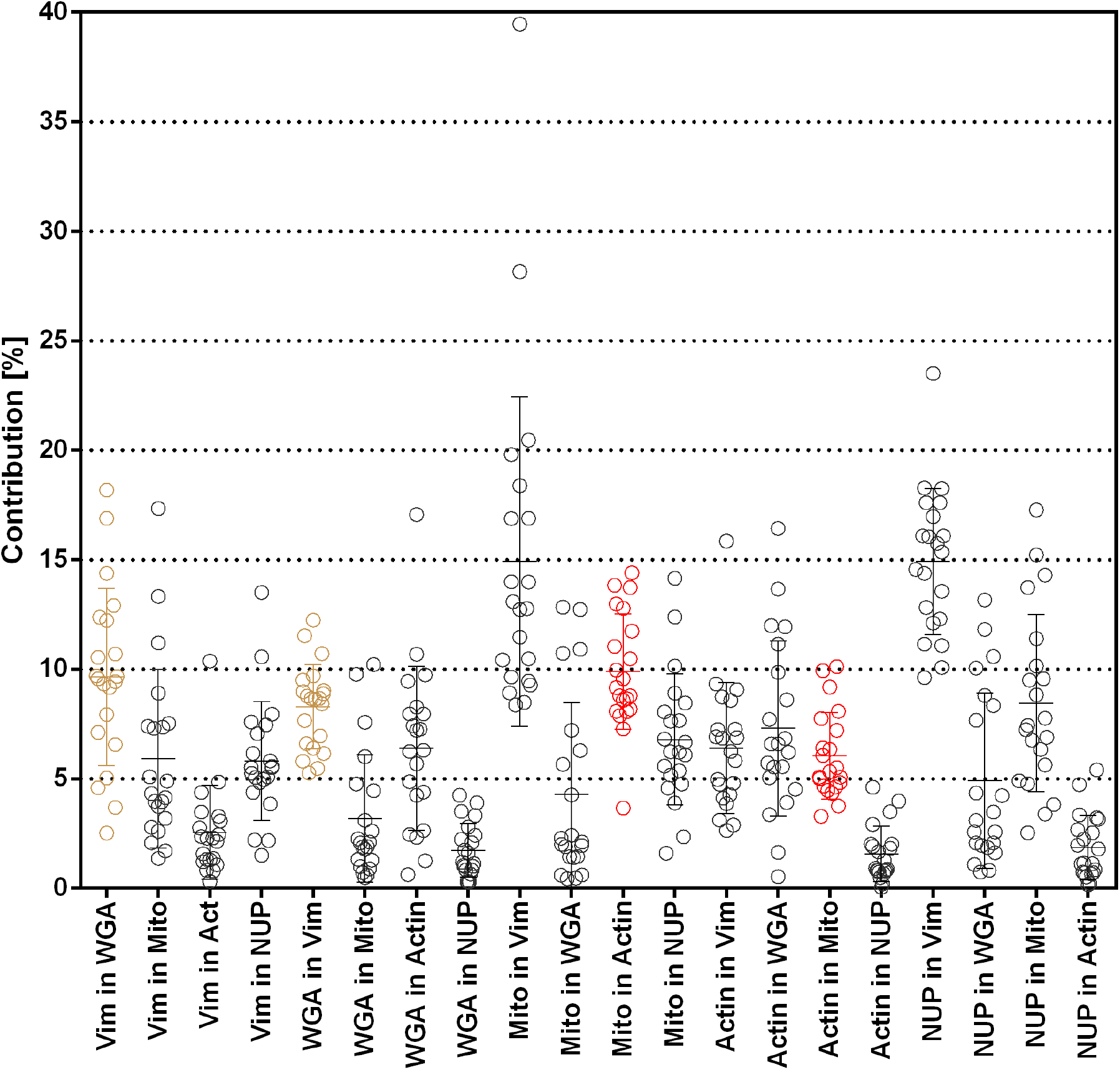
Contribution from channel to channel in five color FLIM STED images, n=21 image sets. Channels generated by phasor separation from the same raw images are shown in the same color, all others in black.

## Discussion

We here show that two fluorochromes per spectral channel can be separated in STED and confocal microscopy if their lifetime is sufficiently different. Exploring this, we generated multi-color FLIM STED images with five colors from three spectral channels, all depleted with 775 nm laser light. We also demonstrate that a fourth spectral channel can be used with 775 nm depletion. Provided two fluorochromes with suitable lifetimes could be identified for each of those spectral channels, eight color STED with a single depletion line would be possible. Long stokes shift dyes with excitation around 500 nm or less, but emission in the red spectral range, might further increase the number of distinguishable labels for a single depletion wavelength.

Already with five colors, we had difficulties to find independent labeling pathways. Typically, labeling of cellular targets occurs with primary antibodies from mouse or rabbit and then secondary antibodies from goat or donkey, which are labeled with the fluorochrome of choice. For some targets, primary antibodies from different animals such as chicken, hamster or rat are available, but these are rare compared to the wealth of rabbit and mouse primaries.

With sufficient budget and experience, primary antibodies could be directly labeled with the desired fluorochrome thus avoiding the problem of limited numbers of species. From a theoretical point of view, this is also desirable since it decreases the distance from the fluorochrome to the actual target, potentially improving the achievable resolution. However, it also decreases signal intensity since several polyclonal secondary antibodies can bind to one primary, multiplying the number of fluorochrome molecules.

Exploiting monoclonal primaries and subtype specific secondary antibodies would increase the number of labeling options. In our study we had to resort to polyclonal commercial secondary antibodies conjugated to fluorochromes, limiting the number of targets we could detect with antibodies from a certain host species to one. We used mouse, rabbit and chicken primaries detected with secondaries from a goat or donkey. These were supplemented with fluorochrome labeled molecules binding directly to cellular targets: WGA-CF594, phalloidin coupled to ATTO 647N or Abberior Star 635P or SPY555-actin.

Apart from the problem of antibody labeling strategies, it is obvious that the fluorochromes that were available to us were not perfect for lifetime separation in STED imaging. Our choices were in particular limited when it came to antibody labels since the two photostable dyes we used in the far red channel, Abberior Star 635P and ATTO647N, have more similar lifetimes when coupled to antibodies. We could separate Alexa Fluor 647 and ATTO 647N antibody labeled signals in STED (unpublished), however, during STED acquisition Alexa Fluor 647 bleached dramatically in our samples.

We hope that the possibilities of multi-color FLIM STED and multi-color FLIM confocal microscopy, together with the advent of respective equipment in many labs, will increase the importance that fluorochrome developers will grant to the lifetime of their dyes and influence future developments accordingly. In general, dyes with a very short lifetime <1 ns will be poor STED dyes, since they will emit a large part of their fluorescence before the depletion effect can take place. Photostable dye pairs with lifetimes above 1 ns and a difference in lifetime of 1 ns or more would be promising candidates, e.g. one lifetime between 1 and 2 ns and one lifetime above 3 ns when coupled to antibodies. Due to our limited resources, we were not able to test further commercially available dyes and it is quite possible that some of them constitute good STED-FLIM pairs when conjugated to antibodies.

For the life science researcher who needs only two or three colors for a STED experiment, it appears to be easiest to use spectrally separable dyes with little spectral crosstalk such as CF680R, Abberior Star 635P and either CF594 or Alexa Fluor 594. SPY555-actin could be used in combination with those fluorochromes. When additional structures need to be labeled, FLIM-STED provides an avenue to achieve this goal.

## Methods

### Cell culture and fluorescence stainings

HeLa-Kyoto cells were kindly provided by Sandra Hake (now Institute for Genetics, Justus-Liebig-Universität Gießen). Cells were passaged twice per week in Dulbecco’s Modified Eagle Medium (DMEM) high glucose, supplemented with Penicillin and Streptomycin and 10% fetal calf serum and grown in an incubator at 37°C with 5%CO_2_. For microscopy, cells were grown on uncoated 15 mm diameter high precision coverslips (170 µm +/-5-10 µm (Marienfeld Superior, Paul Marienfeld GmbH, Lauda-Königshofen,Germany, or Hecht Assistant, Glaswarenfabrik Karl Hecht GmbH, Sondheim vor der Rhön, Germany) in a 10 cm Petri dish for 24 hours.

Live cells staining with wheat germ agglutinin (WGA) CF594 conjugate (Biotium, 29023-1, Stock 1 mg/ml in H2O, diluted 1:500 in medium) was performed for 10 minutes in the incubator. Prestained cells were washed twice in PBS. All specimens were fixed with 4% formaldehyde in PBS made from a methanol-free 16% stock solution (Thermo Fisher Scientific, # 28908). Washing and fixing solutions were prewarmed to 37°C but the 10 min fixation period was at room temperature. Fixed samples were stored in PBS at 4°C until usage.

Permeabilization was on ice with PBS/0.25% Triton X-100 for 6 minutes followed by washing in PBS and incubation for 30 minutes at room temperature in blocking solution, PBS/5% heat inactivated normal goat serum or bovine serum albumin. Primary antibodies were diluted in blocking solution and incubated either over night at 4°C or for 45-60 min at 37°C in a humid chamber. Following three washes with PBS/0.1% Triton X-100, coverslips were rinsed in PBS and incubated with secondary antibodies diluted in blocking solution in a humid chamber for 90 minutes at room temperature or for 45-60 min at 37°C. Three washes in PBS/0.1% Triton X-100 followed. Staining with phalloidin was after a postfixation step with 1% formaldehyde in PBS for 10 min for 45 min at room temperature and followed by another postfixation step, followed by 3 washes with PBS. Coverslips with stained cells were mounted to a glass slide with 7 µl Fluoromount-G (Invitrogen, 00-4958-02). They were sealed with nail polish within 10 minutes after mounting. Air drying of the cells was carefully avoided during the whole procedure.

### Antibodies and fluorophore conjugates

The following antibodies were used at the respective concentrations: Primary: Chicken IgY anti-vimentin polyclonal, Invitrogen, PA1-10003, 1:1000; rabbit anti-TOMM20 polyclonal, Sigma, HPA011562, 1:200; mouse anti-NUP107 monoclonal (39C7); Invitrogen, MA1-10031, 1:100. Secondary: Goat anti-chicken IgY (H+L) cross-adsorbed, Alexa Fluor Plus 594, polyclonal, Invitrogen Thermo Fisher A32759, 1:500 – 1-1000; goat anti-rabbit Abberior STAR 635P, polyclonal, Abberior, 2-0012-007-2, 1:200; Goat anti-rabbit IgG Alexa Fluor Plus 647, polyclonal, Invitrogen Thermo Fisher, A32733, 1:200; Goat anti-mouse IgG ATTO 647N polyclonal, Rockland 610-156-121S, 1:50; donkey anti-mouse IgG CF680R, polyclonal, Sigma-Aldrich, SAB4600207, 1:200. Phalloidin ATTO647N conjugate, ATTO-TEC, AD 647N-81, 1:2000; Abberior STAR 635P phalloidin, Abberior 2-0205-007-0, 20 µg dissolved in 1.77 ml Methanol and diluted 1:400.

### Microscopy

Images were recorded at the Core Facility Bioimaging of the Biomedical center, LMU München. STED was performed on a TCS SP8 WLL STED 3X FALCON from Leica Microsystems CMS, Mannheim, Germany with a 100×1.4 oil immersion “STED white” objective and SMD hybrid photodetectors. The white light laser allowed to pick any wavelength between 470 and 670 nm for excitation. To minimize crosstalk, color channels were recorded in frame sequential mode. A 2D-STED donut was applied. Typical STED images were recorded with 21 - 26 nm pixel size at 400 Hz e.g. for Figure 1 with 1552×1552 pixels with a pixel dwell time of 1.04 µs and a pinhole of 0.93 AU@580 nm. 20 times accumulation for confocal and 100 times accumulation for STED was applied, except for z-stacks (Figure 6) where 40 accumulations were used for STED. Relative laser intensities were typically 9% at 594 nm excitation with 30% STED power, 12% @ 633 with 15% STED, and 10% @ 670 with 10% STED. Recording of a STED image with 1552×1552 pixels and all three spectral ranges with 100 accumulations each took about 13 minutes. No drift was observed during the acquisition.

Samples for which only confocal FLIM images were taken were recorded on a Leica TCS SP8 WLL MP DIVE Falcon (funded by the Deutsche Forschungsgemeinschaft, INST 86/1909-1). For images with 685 nm excitation, a STELLARIS 8 STED FALCON system was kindly made available by Leica Microsystems at their demo labs in Mannheim, Germany.

### Phasor based lifetime separation

Each pixel from the microscopic image is represented by a single dot in the phasor plot, positioned according to its average lifetime. Areas in the plot where dots accumulate are rainbow color coded with blue for low occurrence to red for high accumulation of dots. Lifetime is an intrinsic characteristic of every dye in a given environment, leading to a specific phasor pattern. For an image with two fluorochromes with different lifetimes the phasor plot will typically reveal three scenarios: (i) pixels containing only the shorter lifetime dye, (ii) pixels containing only the longer lifetime dye and (iii) pixels containing a mixture of both. Phasor plot location of pixels containing only one dye will correspond to the location obtained with single stained samples. Pixels with a mixture of both dyes will be located along a line between the patterns of the individual dyes. Their position will depend on the mix of the components from both dyes. This is where the “linear property” of the phasor is powerful: the ratio of the linear combination determines for each image pixel the fraction of the components of the two dyes.

The Leica LAS X software version 4.1.1.23273 (Stellaris) was used for phasor analysis on a separate workstation post acquisition. Using the tools provided in the software, areas identified to contain the phasor pattern of single dyes were selected, and a linear separation was done by the software using the linear property of the phasor plot. A different strategy was applied to images of nuclear pores labeled with CF680R, to suppress ATTO 647N bleed-through, reflection, and the anti-Stokes excitation by the depletion laser. A phasor based filter was applied called Tau STED in the software (see below). This was possible since only one color needed to be extracted. As shown in Figure 1h, a zero intensity ruler of triangular shape was placed both on the reflection (bottom right corner of the phasor plot) and ATTO 647N bleed-through pattern (center left angle) to filter them, to only keep the CF680R signal. Images were subsequently saved and exported. As a result, a separate image for each labeled structure was obtained.

Tau STED applies filtering of pixels based on their phasor signature instead of discarding an arbitrary part of the photons in the whole image as in gated STED. Background from laser reflection and autofluorescence with typically short lifetimes and low resolution photons coming from dyes who emitted before being depleted by the STED laser can be filtered out, while keeping as much information as possible. Within the STED depletion pattern, a diffraction-limited doughnut, intensity is not equally distributed. Since depletion results in a shorter average lifetime of the remaining fluorescence^27^, a certain lifetime distribution is created from the uneven distribution of the depletion laser intensity. This distribution determines the STED trajectory in the phasor plot. Once this STED trajectory is established, it is possible to distinguish photons that are a result of the STED process from any other photon contributions. The photon contributions not correlated to the STED trajectory can be tagged as noise and thus be separated from the signal of interest, as explained above. Based on the characteristics of the fluorochrome undergoing STED, a filter can be applied to single out long lifetime signals within the STED trajectory that more likely originate from the donut center and thereby contribute to a highly resolved image. This solves the problems of reflection and low resolution photons (short lifetimes) and crosstalk (different behavior of different dyes) that can be seen in typical STED multicolor experiments. The accuracy of such determination, as in all lifetime-based approaches, is photon budget dependent.

The low photon numbers typical for STED images are an additional challenge even in the phasor approach. Typical filtering strategies to improve phasor cloud accuracy often fail in these regimes. The use of a complex wavelet filter^28^ both within Tau STED and the phasor separation described above allow to work on the typical STED imaging regimes without a need to further increase the available photons.

No deconvolution was applied, since deconvolution algorithms assume a Gaussian distribution of noise. Such a distribution cannot be assumed as a given, after phasor based separation of fluorochromes. Further resolution increase might be achieved if adapted deconvolution algorithms were available.

Only linear intensity adjustments were applied to images presented in figures.

### Fourier ring correlation

The Fiji-Plugin NanoJ-SQUIRREL^29^ was used for Fourier ring correlation. Complete FLIM-separated images were analyzed, without creating subregions (one block per image). Confocal and STED images were from the exact same sample region, recorded directly after each other and matched in pixel size.

### Crosstalk analysis

This analysis was performed in Fiji^30^ version 1.53f51. For each set of five images generated by phasor separation (or of three images), first a common region for background measurement was identified which contained no structures. The respective average background value was subtracted from each image prior to further analysis to avoid that results would be influenced by different background intensities in different channels.

In multi-color samples it is difficult to accurately quantify crosstalk, since a spatial overlap of differently labeled biological structures, like two crossing filaments, might falsely be interpreted as channel crosstalk. With such samples, it is therefore advisable to focus on areas in which one structure is strongly labeled but no structures of the labels in other channels are present. To achieve this, for each image a high threshold was set such that only intense signals were above it, creating a mask. Often this threshold was near the 97^th^-intensity percentile. Next, a low threshold was determined such that all signals from the structures that were supposed to be in that image were above it. It was typically between the 50 and 70 percentile. All images within a set were compared pairwise: To determine crosstalk from a ‘home channel’ to a ‘neighboring channel’, the high threshold mask from the home channel was taken and the low threshold mask from the neighboring channel was subtracted. For the remaining region of interest (ROI), intensity was measured in both channels and the mean value exported in a csv file.

Bleed-through of the home channel into the neighboring channel was determined in Microsoft Excel as intensity in the ROI of the neighboring channel divided by the intensity in the home channel. To obtain the contribution value, the bleed-through value was normalized by multiplication with the ratio (home/neighbor) of the 99^th^ intensity percentiles of the whole images. A desirable low “contribution” can accordingly be achieved either by low bleed-through or by high intensity of the home channel signal. Graphs, means and standard deviation values were produced in Graphpad Prism 6.01.

## Data availability

Image data will be made available on reasonable request.

## Acknowledgement

We thank Luis Alvarez (Leica Microsystems, Mannheim, Germany) for providing access to a Leica Stellaris system with 685 nm excitation and for helpful discussions. We thank the Deutsche Forschungsgemeinschaft for funding of a Leica SP8 FALCON system (INST 86/1909-1).

## Author contributions

SD conceived the study, designed it with support from MGP and AT, and wrote the manuscript. BB and SR prepared samples and recorded images, supervised by AT, MGP and SD. MGP recorded images and performed the phasor based separation. IN developed the crosstalk-contribution Fiji macros supervised by SD. All authors contributed to the final manuscript.

## Additional information

The authors declare no competing interests.

## Notes

### Competing Interest Statement

The authors have declared no competing interest.

